# Quantifying *Drosophila* Feeding Behavior Using flyPAD and optoPAD

**DOI:** 10.64898/2026.03.20.713238

**Authors:** Nicholas J. Collins, Madison Endres, Irina Sinakevitch, Lisha Shao

## Abstract

Quantifying feeding behavior with high temporal and spatial precision is critical for understanding how internal state, sensory cues, and neural activity shape food intake and dietary choice. Here, we describe a detailed protocol for performing consumption and dietary choice assays in *Drosophila* using the flyPAD/optoPAD system. This method enables simultaneous measurement of feeding events across multiple arenas while allowing precise control of gustatory stimuli and optogenetic stimulation. We provide step-by-step instructions for assay food preparation, flyPAD arena setup, data acquisition, and downstream data organization with suggested analyses. This approach is suitable for studying consumption, nutrient preference, learning, and state-dependent modulation of feeding behaviors, and can be readily adapted for optogenetic manipulations and comparative choice assays.

**Highlights:**

- High throughput, quantitative measurement of *Drosophila* feeding behavior.
- Easy integration of optogenetic stimulation for precise control of neural circuits.
- Compatible with both open-loop and closed-loop optogenetic stimulation paradigms.
- Versatile applications for consumption, nutrient preference, learning, and state-dependent feeding behaviors.
- Step-by-step guidance from arena acquisition and calibration through statistical modeling.

## Before you Begin

## Background

Feeding behavior in *Drosophila* is a widely used and biologically informative variable for understanding how genetic factors, cellular mechanisms, neural circuits, and internal physiological states interact to shape behavior. Alterations in feeding have been used to characterize the effects of social stress^1^, investigate how nutrient valuation contributes to metabolic disease and obesity^2-3^, map associative learning circuits that guide food choice^4-5^, and reveal links between dietary intake, neurodegeneration, and oxidative stress^6^. Because feeding behavior and dietary choice integrate sensory evaluation, motivation, reward, and metabolic demand, disruptions in this behavior often signal underlying changes in neuronal function.

Accordingly, identifying the neural circuits and cellular processes that regulate feeding has become a major research focus. Manipulating these pathways can rescue aberrant feeding phenotypes^7-10^, and offers a tractable window into how physiological state modulates decision-making in the fly brain^11-12^. However, many existing feeding assays lack the temporal precision or neural specificity needed to dissect how circuit activity dynamically shapes feeding decisions. The flyPAD/optoPAD assay addresses these gaps.

As outlined below, flyPAD/optoPAD provides high-resolution detection of feeding on solid food while enabling closed-loop, real-time optogenetic manipulation, making it uniquely suited to reveal how specific neurons influence moment-to-moment feeding behavior.

### Important Definitions

#### flyPAD

For high-resolution, capacitance-based detection of feeding on solid food. Measures total consumption and/or choice between two substrates^13^.

#### optoPAD

Provides the same utility as the flyPAD, with the addition of an optogenetic module for open-loop and closed-loop assays.

#### Open-Loop

A type of optoPAD experiment where optogenetic stimulation is delivered on a continuous, fixed schedule independent of the animal’s behavior.

#### Closed-Loop

A type of optoPAD experiment where optogenetic stimulation is delivered contingent on the animal’s behavior. The flies own behavior facilitates feedback from the optogenetic module^14^.

### Materials and Equipment

**TIMING:** Acquire reagents, consumables, and equipment 1-2 months prior to planned experiments.

Before beginning, the below reagents and consumables (Table 1), as well as equipment (Table 2) should be purchased in advance and set up according to our recommended protocol outlined below. For a summary of equipment and supplies needed to conduct these experiments daily, see Figure 1.

**Table 1.**
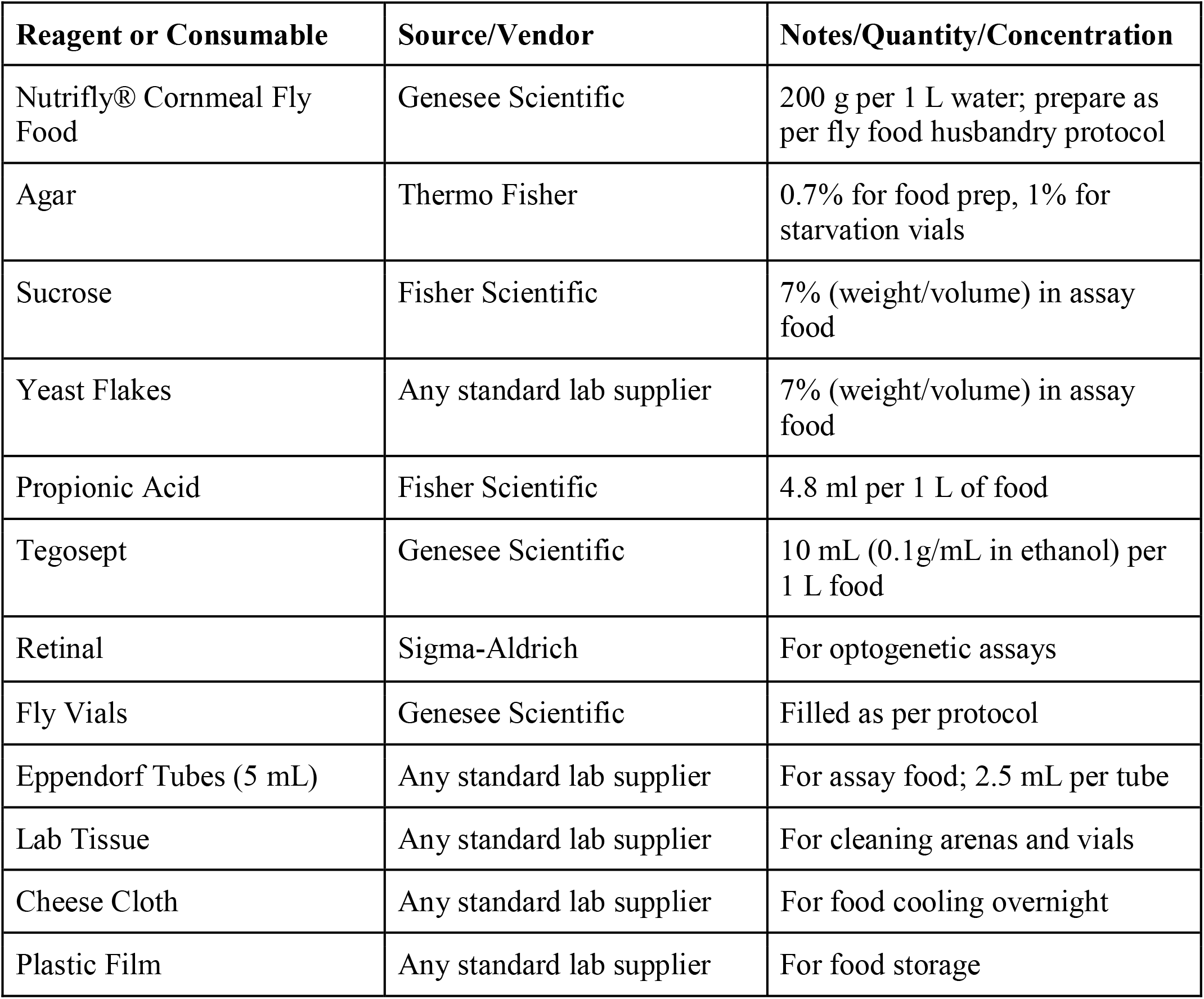
Reagents and Consumables.

| Reagent or Consumable | Source/Vendor | Notes/Quantity/Concentration |
| --- | --- | --- |
| Nutrifly® Cornmeal Fly Food | Genesee Scientific | 200 g per 1 L water; prepare as per fly food husbandry protocol |
| Agar | Thermo Fisher | 0.7% for food prep, 1% for starvation vials |
| Sucrose | Fisher Scientific | 7% (weight/volume) in assay food |
| Yeast Flakes | Any standard lab supplier | 7% (weight/volume) in assay food |
| Propionic Acid | Fisher Scientific | 4.8 ml per 1 L of food |
| Tegosept | Genesee Scientific | 10 mL (0.1 g/mL in ethanol) per 1 L food |
| Retinal | Sigma-Aldrich | For optogenetic assays |
| Fly Vials | Genesee Scientific | Filled as per protocol |
| Eppendorf Tubes (5 mL) | Any standard lab supplier | For assay food; 2.5 mL per tube |
| Lab Tissue | Any standard lab supplier | For cleaning arenas and vials |
| Cheese Cloth | Any standard lab supplier | For food cooling overnight |
| Plastic Film | Any standard lab supplier | For food storage |

**Table 2.**
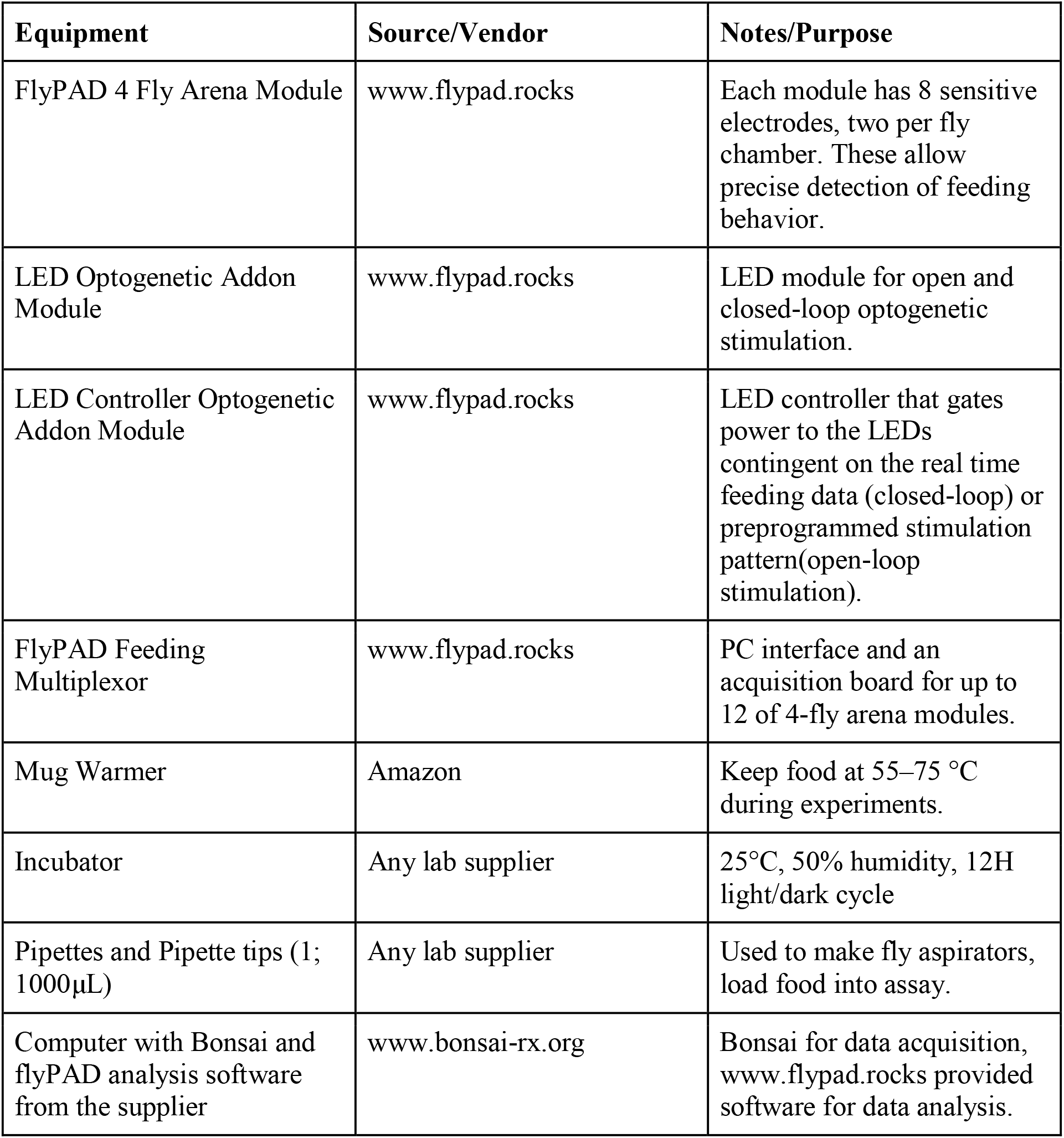
Equipment.

| Equipment | Source/Vendor | Notes/Purpose |
| --- | --- | --- |
| FlyPAD 4 Fly Arena Module | www.flypad.rocks | Each module has 8 sensitive electrodes, two per fly chamber. These allow precise detection of feeding behavior. |
| LED Optogenetic Addon Module | www.flypad.rocks | LED module for open and closed-loop optogenetic stimulation. |
| LED Controller Optogenetic Addon Module | www.flypad.rocks | LED controller that gates power to the LEDs contingent on the real time feeding data (closed-loop) or preprogrammed stimulation pattern(open-loop stimulation). |
| FlyPAD Feeding Multiplexor | www.flypad.rocks | PC interface and an acquisition board for up to 12 of 4-fly arena modules. |
| Mug Warmer | Amazon | Keep food at 55–75 °C during experiments. |
| Incubator | Any lab supplier | 25°C, 50% humidity, 12H light/dark cycle |
| Pipettes and Pipette tips (1; 1000µL) | Any lab supplier | Used to make fly aspirators, load food into assay. |
| Computer with Bonsai and flyPAD analysis software from the supplier | www.bonsai-rx.org | Bonsai for data acquisition, www.flypad.rocks provided software for data analysis. |

**Figure 1.**
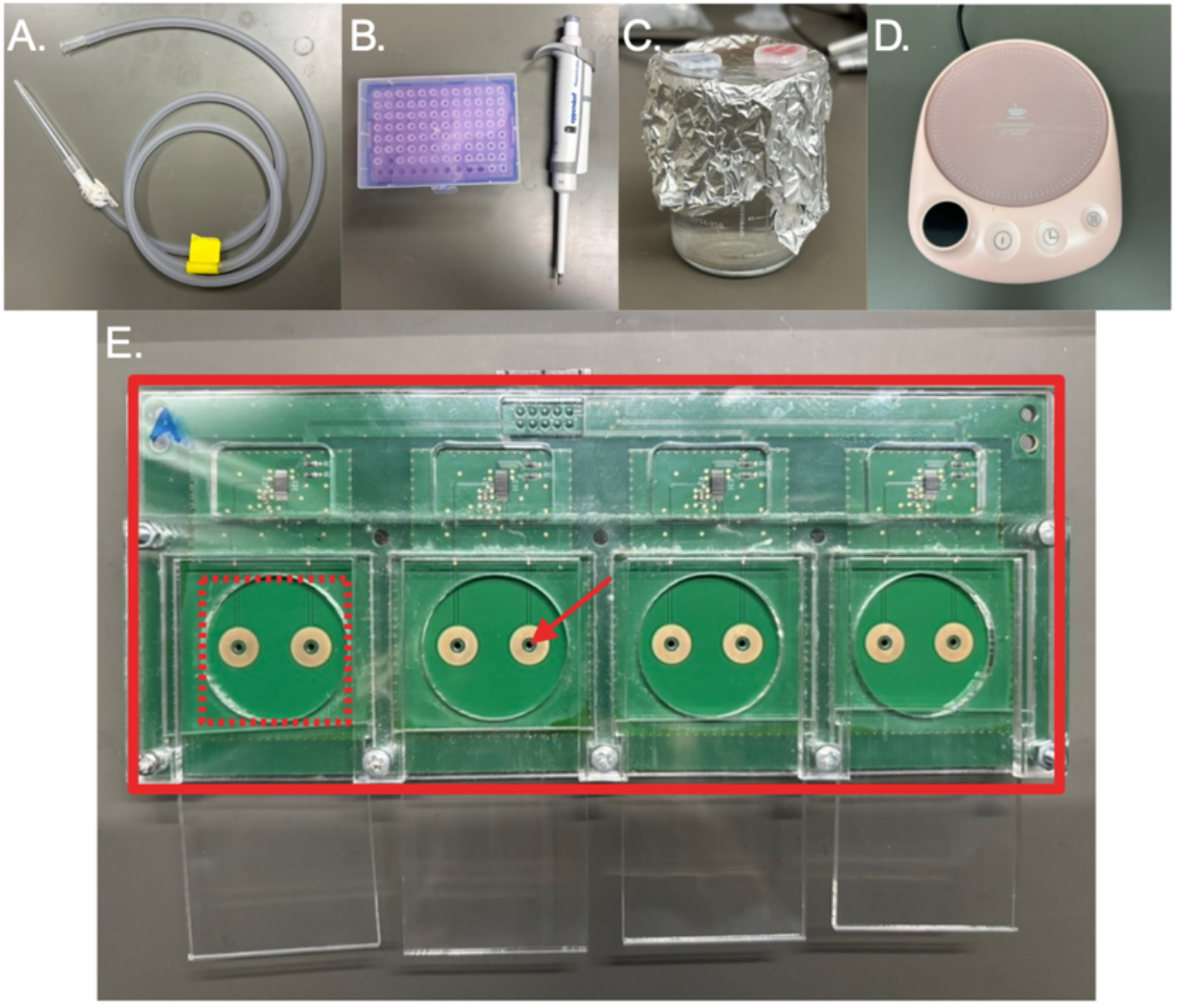
Required materials for the flyPAD assay. (A) A fly aspirator is used to transfer a single fly into a flyPAD chamber. The aspirator is constructed from plastic tubing, cheese cloth, and a pipette tip. (B) A 10μL pipette and tips are used to load the appropriate food media into each flyPAD channel (red arrow). Approximately 3μL of media is loaded per channel. (C) Food media are stored in 5 mL Eppendorf tubes prior to heating. They are then heated and maintained in a hot water bath during the loading process. (D) A standard mug warmer is used to keep the food media in a liquid state between experiments, reducing the need for repeated reheating. (E) The flyPAD unit (solid red rectangle) is used for experimentation. One fly is loaded into each flyPAD chamber (dashed red rectangle). The transparent top panel of each chamber can slide open to allow loading of food media and flies during assay preparation.

### Setting up and procurement of the flyPAD Units and optoPAD LED modules

**TIMING:** 1-2 months prior to start of experiment

This section describes acquisition, initial setup, and pre-experimental validation of the flyPAD/optoPAD system. Early setup is critical to allow sufficient time for hardware troubleshooting, software configuration, and pilot testing prior to data collection.

1. Procurement
  a. Obtain flyPAD units, sensor boards, control electronics, optogenetic LED modules, power supplies, and all required cables from the manufacturer.
  b. Confirm with manufacturer that purchased units are compatible with both open-loop and closed-loop optogenetic configurations if desired.
  c. Ensure that replacement units, covers, screws, and cables are available prior to beginning experiments.
2. Initial Hardware setup
  a. Place the flyPAD unit on a vibration-minimized, level benchtop in a temperature- and humidity-controlled behavior testing room.
  b. Connect flyPAD sensor boards and optoPAD optogenetic control modules to the acquisition computer according to manufacturer specifications.
  c. Secure flyPAD units evenly to sensor boards, ensuring that all contact points are flush and screws are tightened uniformly to prevent signal drift.
3. Software installation and configuration
  a. Install Bonsai (behavioral acquisition software) and required flyPAD/optoPAD packages on the acquisition computer. **NOTE:** The correct version of Bonsai to install is provided by the manufacturer.
  b. Verify correct COM port assignments for flyPAD sensors and optoPAD controllers (port numbers may vary between systems).
  c. Load open-loop and closed-loop Bonsai workflows and confirm that all nodes execute without errors.
4. Pre-experimental calibration and validation
  a. Run a baseline channel test with empty arenas to confirm stable capacitance signals across all channels (Figure 2).
  b. Load agar-only food (1% agar in distilled water) into test wells and initiate a short test recording to verify that all channels show dynamic, non-flat traces (see Figure 2 and Figure 3).
    i. Click on “Start”
    ii. Double click on “flyPAD sensors data”.
    iii. Verify all lines are in a zig-zag fashion.
    iv. If lines are straight, seek troubleshooting.
5. Temperature and Humidity
  a. Maintain testing room temperature at 22–25°C and relative humidity between 40-55%, as environmental variability can affect both feeding behavior and sensor sensitivity.
  b. Conduct experiments in darkness or under controlled lighting conditions to prevent unintended visual stimulation.

**Figure 2.**
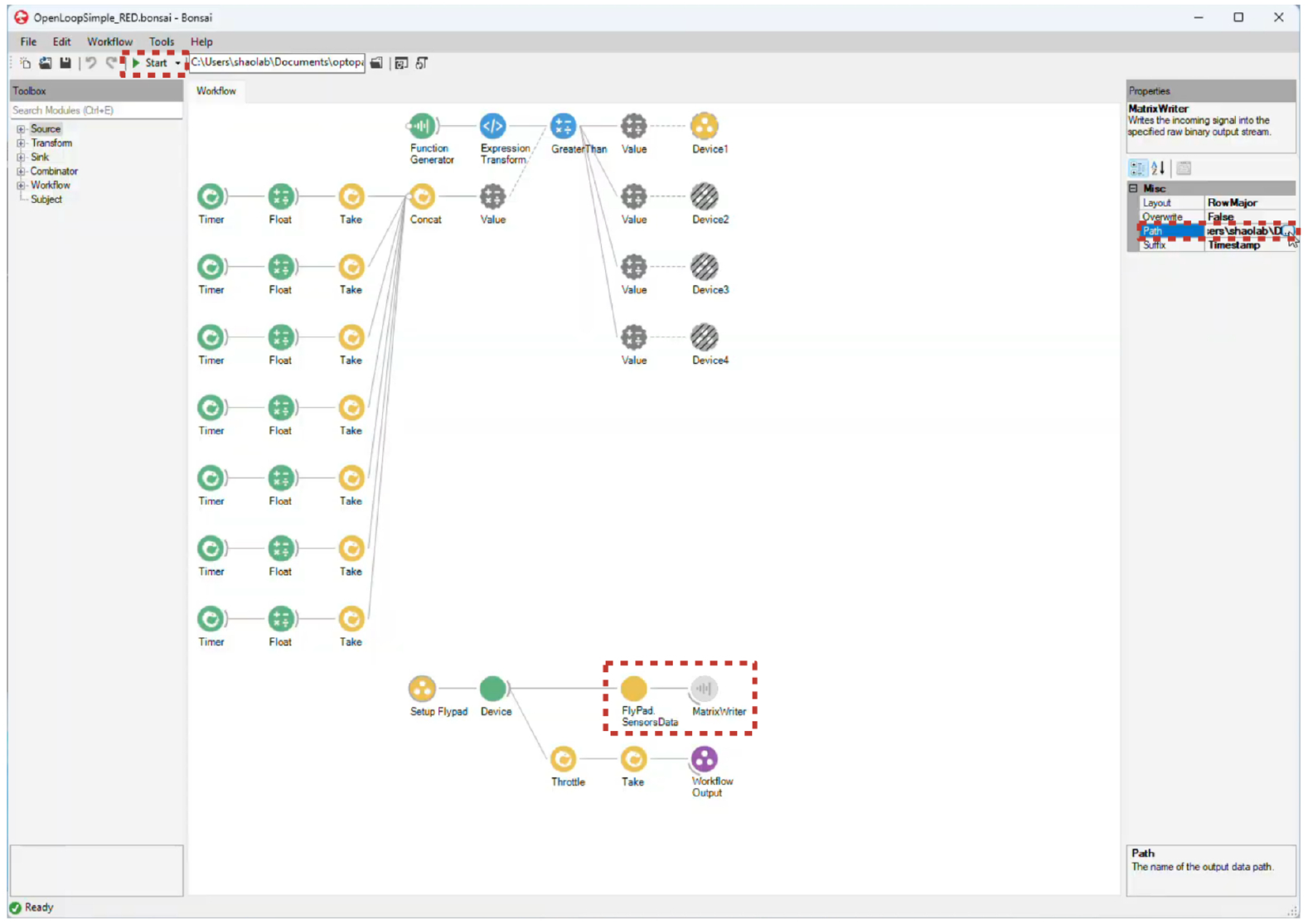
Key features of the Bonsai setup and assay initiation. Shown is the typical appearance of a Bonsai workflow file. Critical parameters that should be edited prior to assay initiation are indicated by dashed boxes. MatrixWriter” specifies where the Bonsai output file will be saved once the assay is stopped, and the “Path” field can be modified to direct the file to the appropriate storage location. “flyPADSensorsData” should be opened during media loading to confirm that all channels are functioning properly. After these settings have been configured for the experiment, the “Start” button can be clicked to initiate the assay.

**Figure 3.**
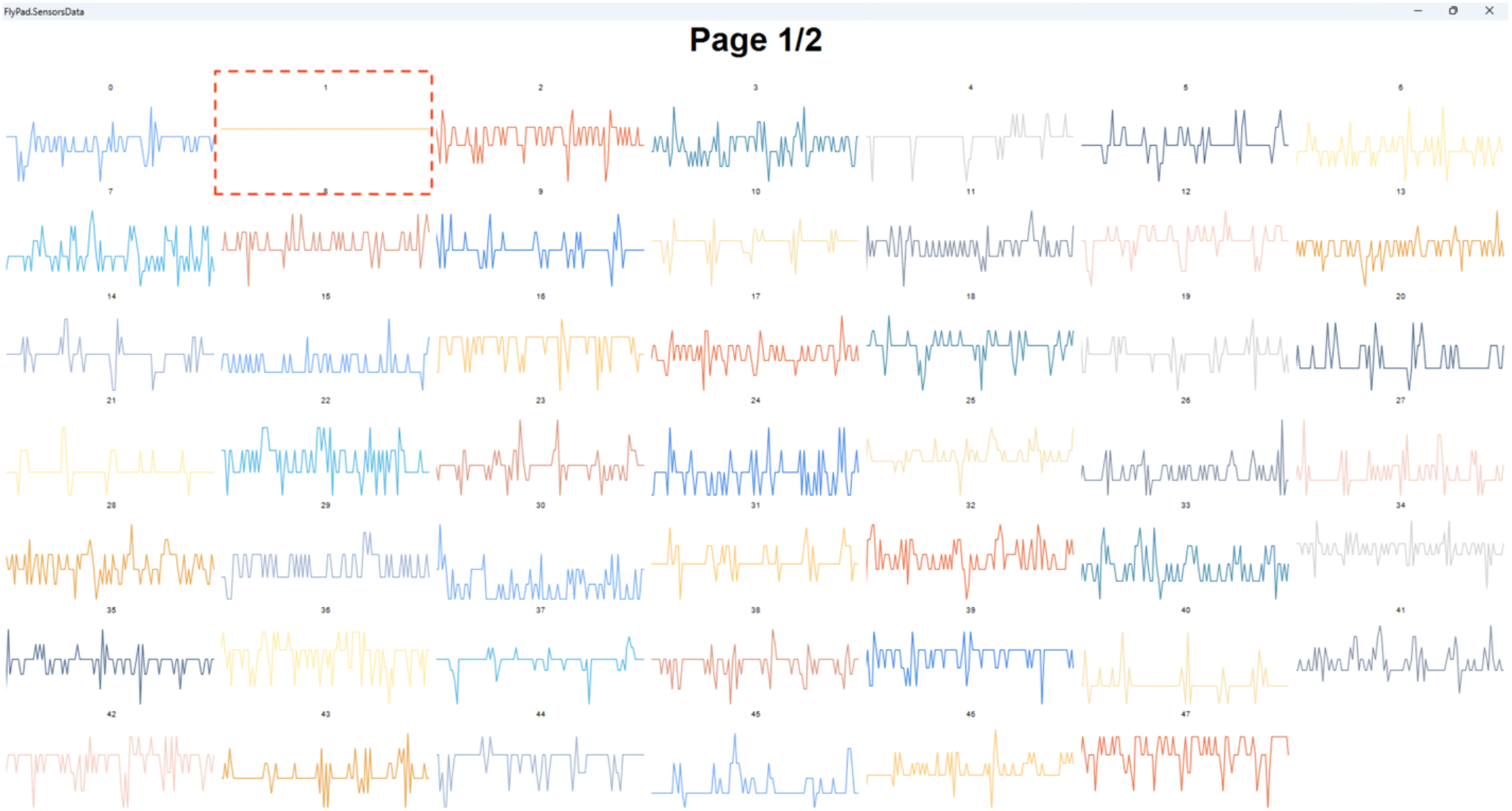
Example of flyPAD sensor data. During loading of food media into the flyPAD channels, sensor data should be monitored to verify proper loading. Each panel corresponds to an individual flyPAD channel within a chamber and is numbered accordingly. After successful media loading, sensors should display a characteristic zig-zag pattern, indicating that sensor function has not been compromised by the media. In contrast, a flat signal (dashed box) suggests that media has spilled onto the sensor, which can interfere with subsequent data analysis. To avoid this issue, channels should be cleaned thoroughly and media loaded carefully. Once all sensors exhibit the zig-zag pattern, media loading is complete and the assay can be initiated.

**CRITICAL:** Allocate sufficient time for pilot experiments to confirm reliable feeding engagement, sensor stability, and accurate synchronization of optogenetic stimulation before collecting experimental data.

### Cooking, dispensing, and storing standard food for husbandry

**TIMING:** 3-4 weeks prior to experiment; 1-2 hours.

1. Cooking fly food for husbandry (Base volume: 1L). Scale as necessary.
  a. Add 1L of distilled water and 200g of Nutri-Fly® to a large pot.
  b. Place pot on an electric stove top, set on high heat (simmer setting, approximately between 85-96□. Stir continuously to avoid food sticking to the bottom of the pot.
  c. Once the mixture reaches boiling point, put the burner to a higher heat (approximately 100□), and continuously stir for an additional 10-15 minutes.
  d. Take off the food from the heat. Turn off the heat, and allow the food to cool to 70□ with a continuous stir.
  e. Add propionic acid (4.8 mL) and Tegosept (10 mL) as preservatives. Stir food to evenly distribute ingredients.
2. Dispensing
  a. It is recommended to use a rapid dispensing system (see equipment below)
    i. For husbandry vials, fill to ∼2-3cm depth.
    ii. For collection vials (for collecting flies prior to beginning assay) fill to ∼1-2cm.
3. Storage
  a. Cover newly created food vials with cheese cloth and allow food to solidify overnight (6-8H).
  b. Food can then be stored in 4□for approximately 1 month. **NOTE:** Do NOT insert vial plugs during cooling. **NOTE**: Cover tightly with plastic film and cover in a plastic bag.

### Retinal Food for husbandry and optoPAD

**TIMING:** ∼2 weeks prior to experiment

1. Cooking Retinal food (optogenetic experiments)
  a. Prepare retinal stock solution
    i. Dissolve all-trans retinal in ethanol to make a 100mM stock solution.
  b. Follow the standard food preparation steps outlined above.
2. Addition of Retinal to standard food
  a. Once food reaches a temperature of 60□, turn off all of the lights in the laboratory. Dim ambient lighting may be used.
  b. Add the appropriate amount of retinal.
    i. Husbandry vials: 1 mL retinal stock solution per 500 mL food.
    ii. Collection vials: 1 mL retinal stock solution per 250 mL food.
3. Repeat standard food protocol for cooling, dispensing, and storage.

**CRITICAL 1:** Place retinal food into a light-sensitive plastic bag to prevent degradation.

**CRITICAL 2:** You may prepare 0.2 mM retinal-containing food for husbandry (1:500 ratio). This is recommended if flies will be collected from these vials and immediately used in behavioral experiments. For collection vials, you may use 0.4 mM retinal containing food (1:250 ratio), and flies should remain on this food for at least 3 days prior to the experiment.

### Preparing Experimental Flies

**TIMING:** ∼2 weeks prior to experiment

1. Maintain newly acquired stocks (e.g., from the Bloomington Stock Center) in husbandry vials as described above in a 25□ incubator.
  a. Temperatures between 18□ to 27□ can be used to slow or speed up development, respectively.
  b. Fly incubators should be set to allow for a 12H dark/light cycle.
  c. Humidity should be set to 50-55%.
2. Transfer flies to a new vial with food every 3-5 days, to prevent overcrowding and to easily identify virgin progeny (if desired).
3. Flies in the assay may be:
  a. Control strains (*Canton-S, w1118*, etc.) or
  b. Progeny from Gal4/UAS crosses, including crosses for optogenetic experiments (see protocol below).
4. Ideal sample collection conditions:
  a. 20-25 flies per group is often sufficient sample size for most applications of the flyPAD/optoPAD assay.
    i. Run a power analysis if desired to confirm adequate sample size.
  b. Collected flies can be kept in collection vials prior to beginning the experiment.
  c. For most experiments, flies should be between 3-7 days old.

**CRITICAL:** Do NOT use CO_2_ within 24h of beginning the flyPAD experiment, as this can influence fly behavior. Instead, use a fly aspirator to transfer flies into the appropriate arenas.

### Preparing Experimental Food for flyPAD and/or optoPAD Assays

**TIMING:** 1 week prior to experiment; 15 minutes.

The flyPAD/optoPAD assay allows the experimenter to choose any two food types for presentation to the fly. Each food type used in our lab, including its specific ingredients, preparation instructions, and storage conditions, is described individually below. Sucrose-based food is provided as the primary example. Any deviations from this procedure (steps omitted or added) for other food types are indicated in Table 3.

**Table 3.**
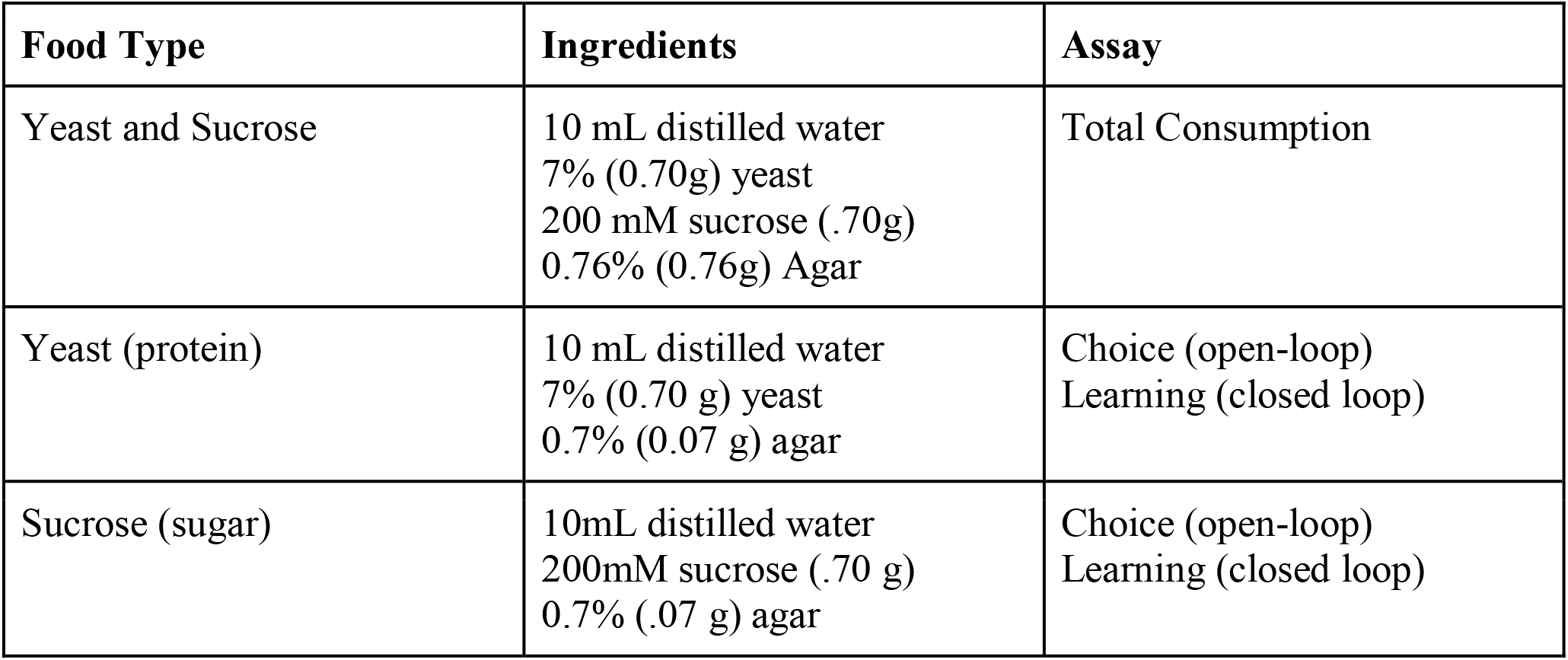
Experimental flyPAD/OptoPad Food Recipes.

**Table 4:**
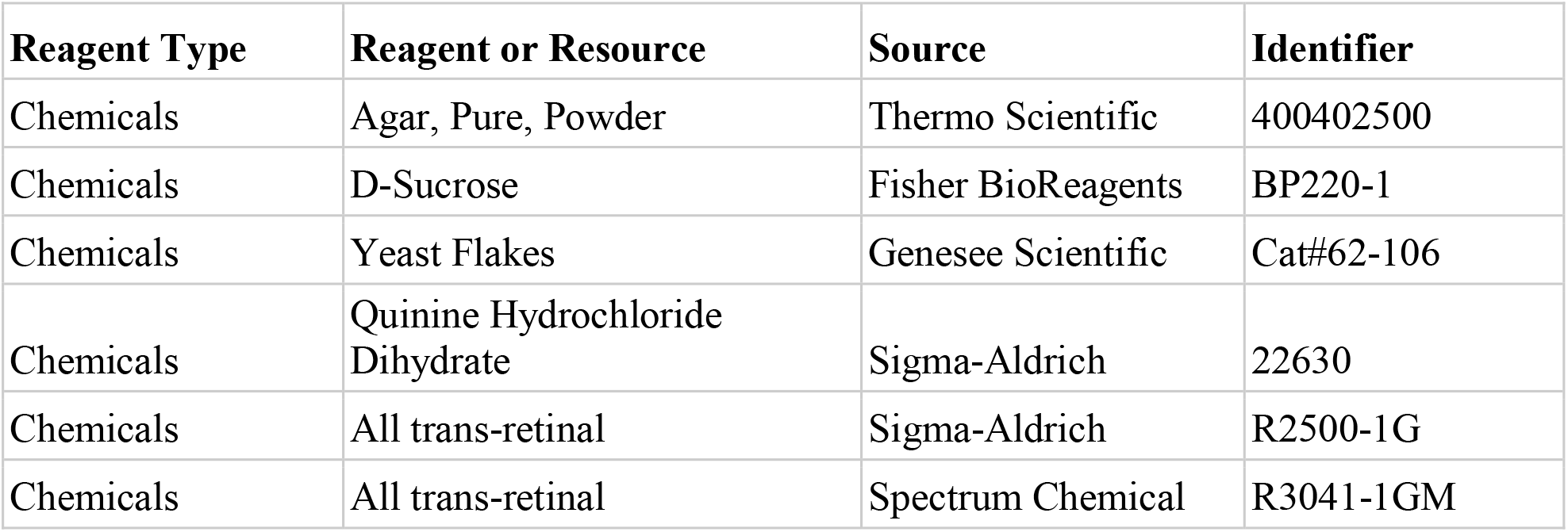
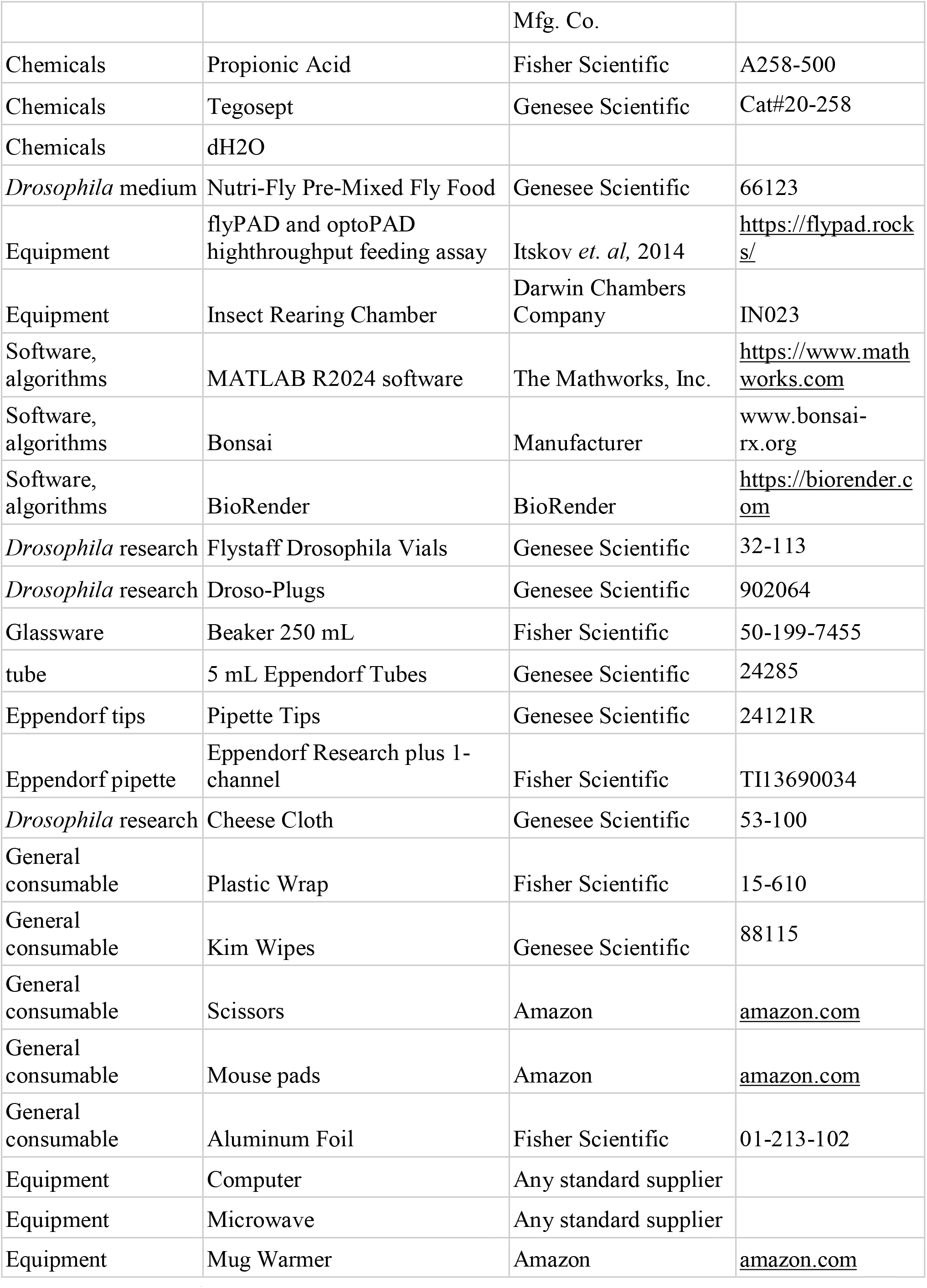
Key Resources.

1. Preparation of sucrose food
  a. To a microwave-safe beaker, add 10 mL of hot distilled water (up to 95□), 0.70g sucrose (200mM) and 0.07g Agar (0.7%). For consistency, place the mixture on the mug warmer on 55-75□, stirring every 15s until all components are completely dissolved.
  b. Pour sucrose food into a 5 mL Eppendorf tube, filled approximately half-way (approximately 2.5 mL).
  c. This will create 4 tubes of food, which ∼8 experiments.
  d. Ensure food solidifies within 1 hour at RT.
2. Food can be stored at RT for approximately 2 weeks or reheated twice.

### Innovation

The flyPAD/optoPAD feeding assay provides several advantages over existing feeding paradigms such as Con-Ex (blue dye), CAFE, and FLIC^13,15-17^, flyPAD/optoPAD combines high-resolution, capacitance-based detection of feeding on solid food with closed-loop, millisecond-precision optogenetic control, enabling neural circuit perturbations that occur strictly during active feeding bouts. Compare to Con-Ex, which provides accurate but low-resolution measures of total consumption, or FLIC, which offers high-resolution detection of liquid-food interactions, the flyPAD/optoPAD systems allows researchers to causally link neuronal activity to microstructural feeding decisions in real time on solid food.

This integration of precise behavioral readout and temporally locked circuit manipulation makes the flyPAD/optoPAD system uniquely suited for dissecting how specific neurons drive feeding preference, consummatory microstructure, and the moment-to-moment dynamics of ingestive behavior.

### Limitations

While the flyPAD/optoPAD offers several advantages, there are also some limitations worth noting. First, the assay only allows for the choice between two different food sources, which may not be representative and engage the full behavioral repertoire. Second, the chambers are only in theory designed to study the behavior of individual flies; it is not practical to use the device to study feeding behavior in group dynamics. Third, the assay is mainly optimized for studying short-term feeding behavior within an hour. As such, it is critical the experimenter considers the importance of circadian rhythms, which are known to influence overall consumption patterns in *Drosophila*^18^.

### Institutional Permissions

There are no specific institutional permissions required, other than standard bio-safety training as directed by local and federal requirements.

### Preparing data files prior to assay

1. Establishing a directory for saved data
  a. Open the “Bonsai” software on the acquisition PC.
  b. Choose the relevant protocol you wish to run (provided from manufacturer).
    i. Ex.: “OpenLoopSimple”
  c. Double click on “matrix-writer”
    i. A panel should open up to the right on the screen.
  d. Where it says “path”, select the ellipsis.
  e. Select the directory where you want your data file to be saved.
  **NOTE**: It is recommended to save each experiment in its own folder.
2. Naming the File (see Figure 4)
  a. Files will be named according to experimental conditions (number of groups and number of flies per group).
    i. For example, C01_01_08_C02_09_16
    ii. This would suggest that you have two groups (C01 and C02).
    iii. Each one of those groups will utilize 4 chambers (01_08 refers to EACH flyPAD channels, so 01 and 02 combined make up the first chamber).
3. Preparing a text/log file for metadata
  a. In the experimental folder, make a.txt file
  b. Each group will be labeled using the following convention:
    i. Events.ConditionLabel{1} = ‘Group 1 information’
    ii. Events.ConditionLabel{2} = ‘Group 2 information’
  c. Each substrate (food) will be labeled with the following convention:
    i. Events.SubstrateLabel{1} = ‘food pipetted on the LEFT in each chamber’
    ii. Events.SubstrateLabel{2}= ‘food pipetted on the RIGHT in each chamber’

**Figure 4.**
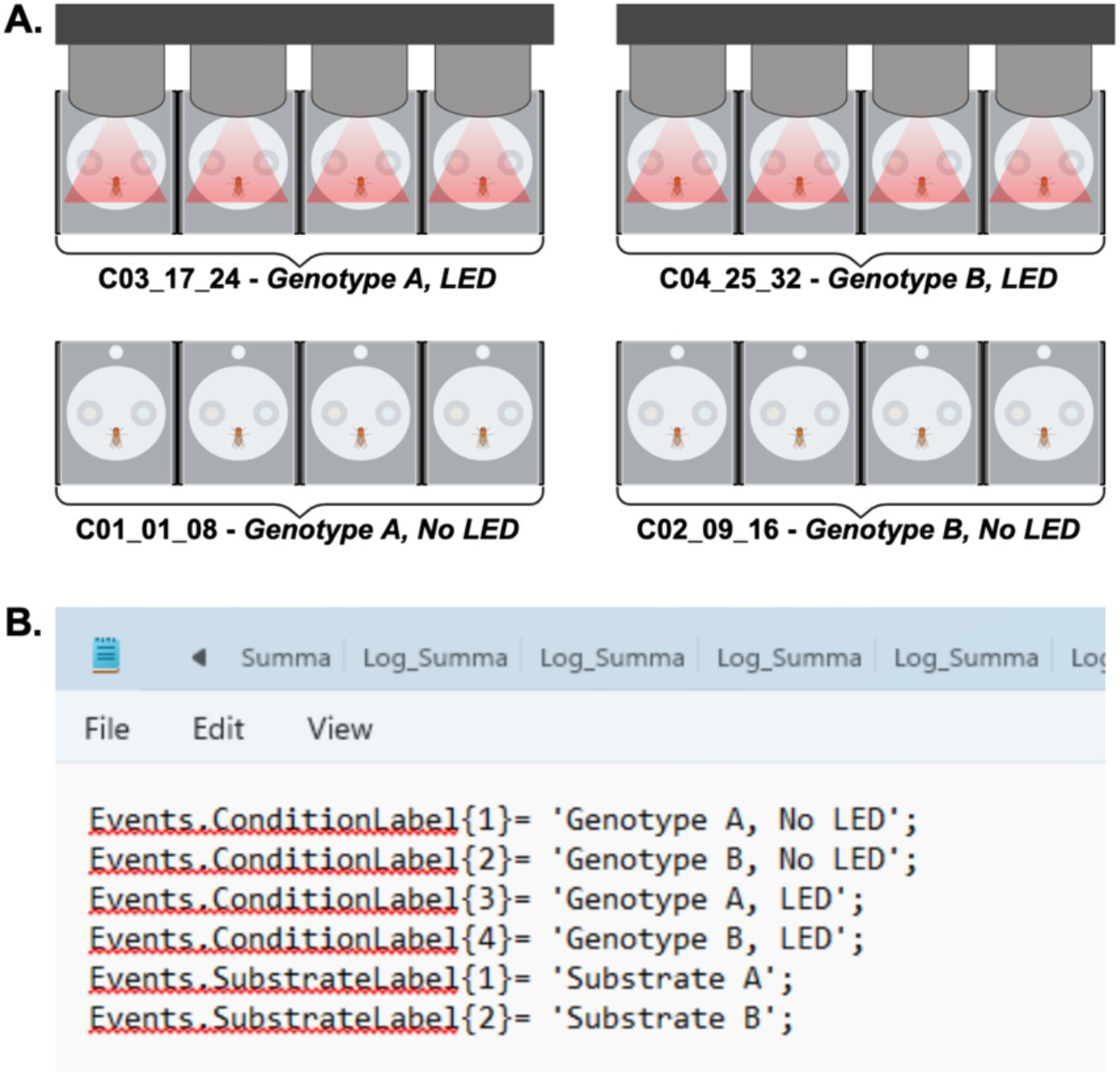
Example of Fly/optoPAD choice assay setup and corresponding log file. (A) Schematic of a flyPad experiment with four experimental conditions looking at two different genotypes (Genotype A or B) and two illumination states (LED on or off). Each flyPAD channel contains one of two food substrates (e.g. yeast on the right, sucrose on the left) with one fly loaded per chamber. Continuous LED illumination (e.g. optogenetic stimulation) is applied to the top row of chambers, while the bottom row remains unilluminated. Each genotype is tested under both LED conditions (example shown: n=4 per condition). Condition labels (C0#) correspond to entries in the Bonsai and log files. The numeric ranges (e.g. 01_08) indicate the channel numbers assigned to each condition. Channels are numbered from left to right, starting with 01 at the bottom left. (B) Example log file corresponding to the experimental setup in (A) to be used as a reference for downstream MATLAB analysis. Events.ConditionLabel {#} specifies the genotype and LED condition for each group. This may also include additional factors such as: fly sex, mating status, starvation condition, etc. The number of condition labels in the log file must match those defined in the Bonsai file. Events.SubstrateLabel {1} and {2} indicate the substrates loaded on the left and right channels of each chamber, respectively. If only one substrate is tested, only SubstrateLabel {1} is required.

**CRITICAL 1:**. How you label the data file indicates where each group is located and what conditions they are exposed to for the analysis. However, files can be renamed after the experiment is run.

**CRITICAL 2:** Be sure to remember and log which food of interest is pipetted where and keep this consistent *within* each experiment. You may alternate food sides *between* experiments.

### Behavior Protocol: flyPAD assay

The flyPAD experiment allows the fly openly feeding between two choice points. This type of experiment is ideal for studying:

a. Total consumption: baseline feeding behavior on a standard food.
b. Choice: baseline choice feeding behavior between two different types of food.

For example, you may wish to determine the total amount of food a particular fly line consumes over the 50-minute testing period (Figure 5), or if the fly prefers protein vs sucrose (Figure 6)

**Figure 5.**
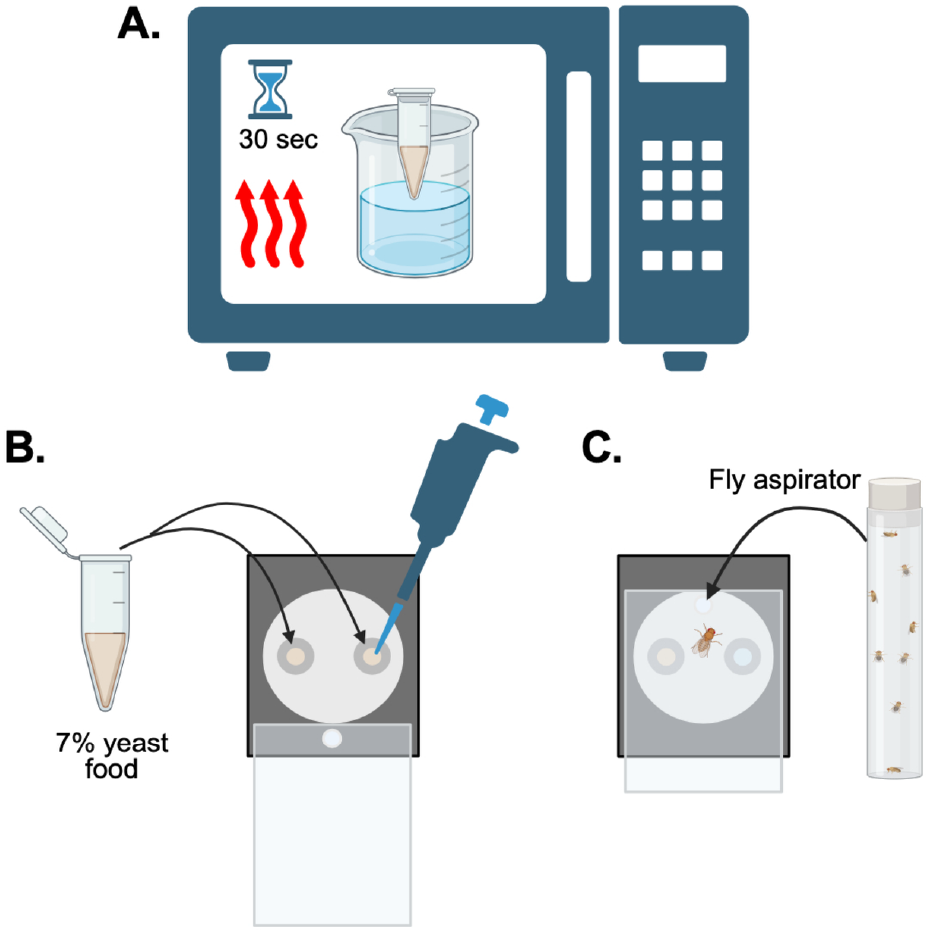
General procedure for open-loop flyPAD assay to measure total consumption. (A) Food media is liquified by microwaving for ∼30s in 10s intervals. The media should not reach a boiling state. The media is then maintained in a hot water bath to prevent solidification prior to loading. (B) Approximately 3μL of food media is pipetted into each flyPAD channel, taking on a dome-like shape. For total consumption measurements, identical media is loaded into both channels. (C) A single fly is loaded into each flyPAD chamber using a fly aspirator through a small opening in the transparent top panel. The chamber is then fully sealed, and the assay can begin.

**Figure 6.**
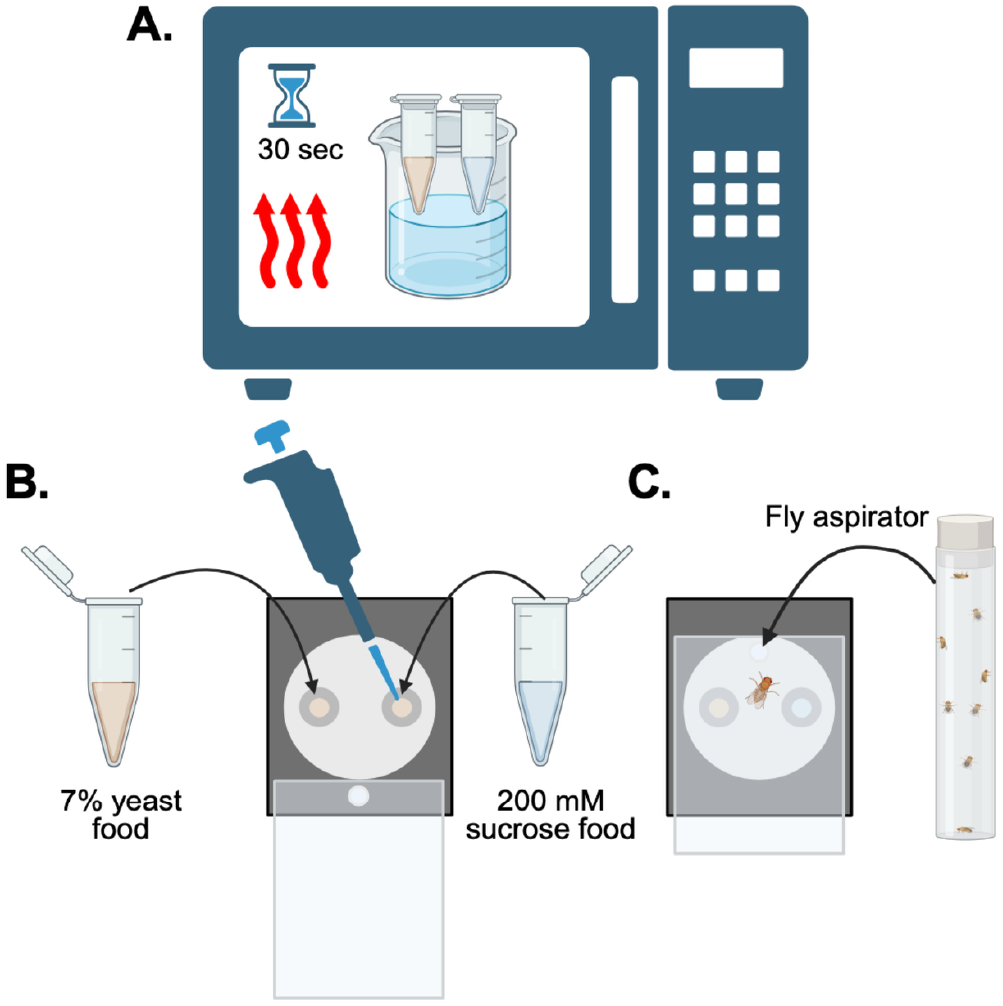
General procedure for open loop flyPAD assay to measure two-choice. (A) Each food media is liquified by microwaving for ∼30s in 10s intervals. The media should not reach a boiling state. The media are then maintained in a hot water bath to prevent solidification prior to loading. (B) Approximately 2-3μL of food media is pipetted into each flyPAD channel, taking on a dome-like shape. For choice experiments, one media type should be consistently loaded on the left channel, and the other media on the right channel. (C) A single fly is loaded into each flyPAD chamber using a fly aspirator through a small opening in the transparent top panel. The chamber is then fully sealed, and the assay can begin.

1. Pre-Experiment PC preparation
  a. Turn on the PC that is associated with your flyPAD rig.
  b. Open up the “Bonsai” software.
  c. Select your protocol of choice, in this case choose an open-loop protocol.
  d. Create a directory and.txt file as directed above.
2. Warming food for the assay
  a. Place 5 mL tubes of sucrose and protein and on a microwave-safe tray, with the lids open.
  b. Fill a beaker (200 mL) with ∼150 mL of water.
  c. Microwave the tubes and the beaker for approximately 1 minute.
    i. While microwaving check every 15-20 seconds.
    ii. Sucrose and protein will be done when they are in liquid form and warm to the touch.
  d. Place tin foil over the water-filled beaker and push the tubes with your food through to the other side. Store tubes in warm water (see Figure 1).
    i. Place the beaker with tubes on the mug warmer at 55 □ during experiments.
    ii. This is to keep the food warm and liquified while you are working with it.
  **CRITICAL:** You do NOT want the food to solidify too early in the paradigm, as it makes it difficult for the flies to consume. Ensure that the food is indeed warm, in a liquid state, and the water remains warm to the touch while working with it.
3. Loading food into flyPAD channels (Substrate A; Left Channel)
  a. Set a 10 μL pipette to 10 μL.
  b. Load desired food into the LEFT channel within each arena.
    i. Each channel doesn’t get a precise amount of food. Your goal here is to provide a base layer of food.
    ii. We recommend ∼3.33 μL into each (3 chambers per pipette full).
  **CRITICAL**: While pipetting the food, you should NOT be getting any food on the brass section of the arena, as this will interfere with accuracy of results and require more troubleshooting (see below).
4. Loading food into flyPAD channels (Substrate B; Right Channel, see Figure 5).
  a. Repeat step 3 for your alternate (two-choice assay) or same (consumption assay) substrate for the right channel.
  **NOTE**: If running a total-consumption experiment, you should load the same substrate in BOTH channels (Figure 6). **CRITICAL 1**: The sucrose food is transparent, so avoiding the brass becomes more difficult. As such, you may put a 1:200 ratio of blue dye into the sucrose food if necessary. Our lab has determined no significant differences with fly feeding between sucrose or sucrose with dye containing foods (data unpublished but available upon request). **CRITICAL 2:** Please ensure you are changing your pipette tip after different foods to prevent cross-contamination.
5. Add an additional layer of food of each substrate to create a dome shape.
  a. Pipette 1.25μL of substrate A and B over each (respectively).
  b. See Figure 7 for an illustrative example.
6. Pre-Experimental Validation
  a. See step 4 of the setting up and procurement of the flyPAD Units and optoPAD LED modules section.
  b. Once you confirm all capacitance lines are moving and wavy, select “stop” on the bonsai software.
    i. If any are flat, wipe up excess food using a lint-free cloth and re-check.

**Figure 7.**
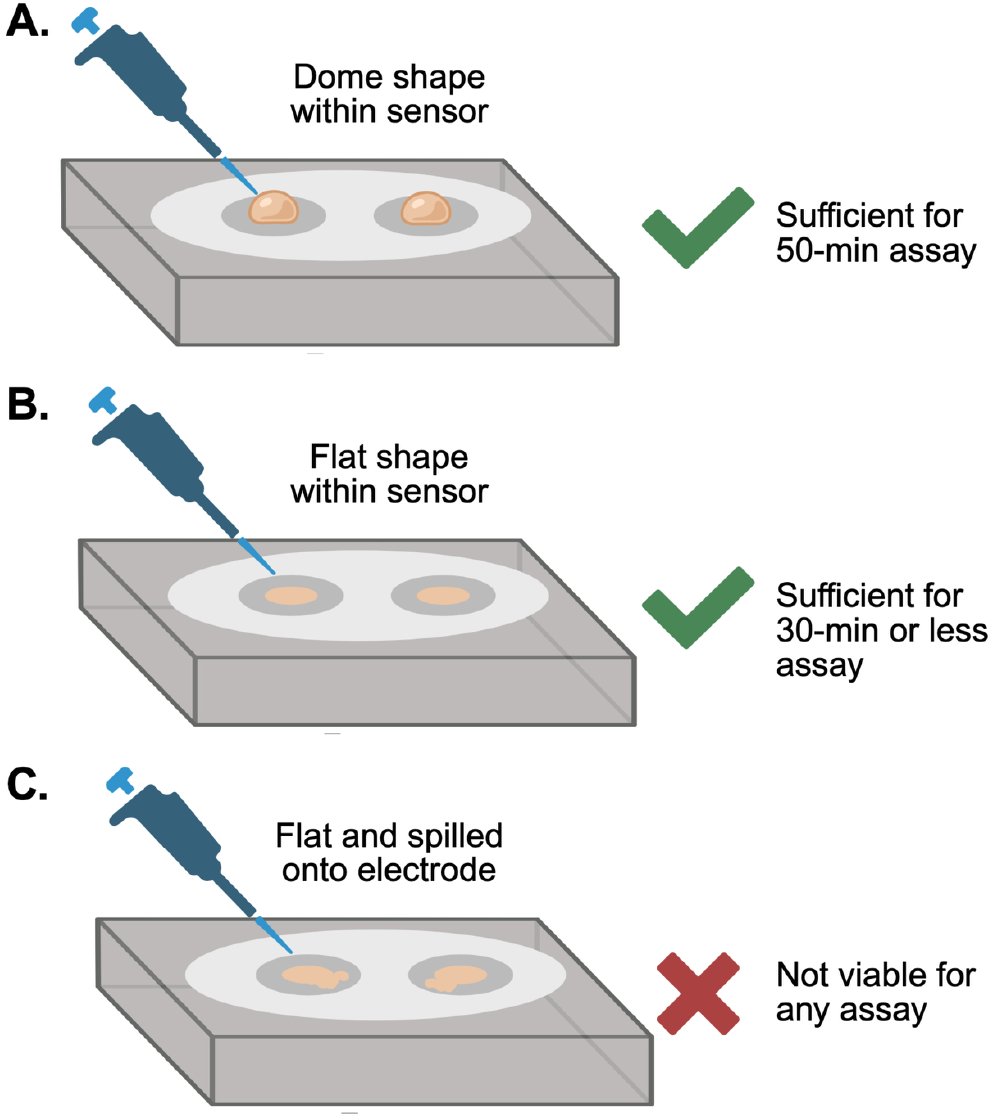
Proper loading of food media into assay chambers. Using a micropipette, dispense ∼2-3μL of liquified food media into the center of each channel. In longer assays spanning 50 minutes, the media should form a raised, dome-like meniscus (A), which enables effective interaction with the food by the fly. In shorter assays spanning 30 minutes or less, food can take on a flat shape within the channel, without spilling onto the sensor (B). If the media spreads into a flat layer that spills onto the electrode (C), the data will not be viable for any assay. If this occurs, wipe the sensor with a lab tissue and add ∼1μL of additional media to the top until a domed shape is achieved.

**Figure 8.**
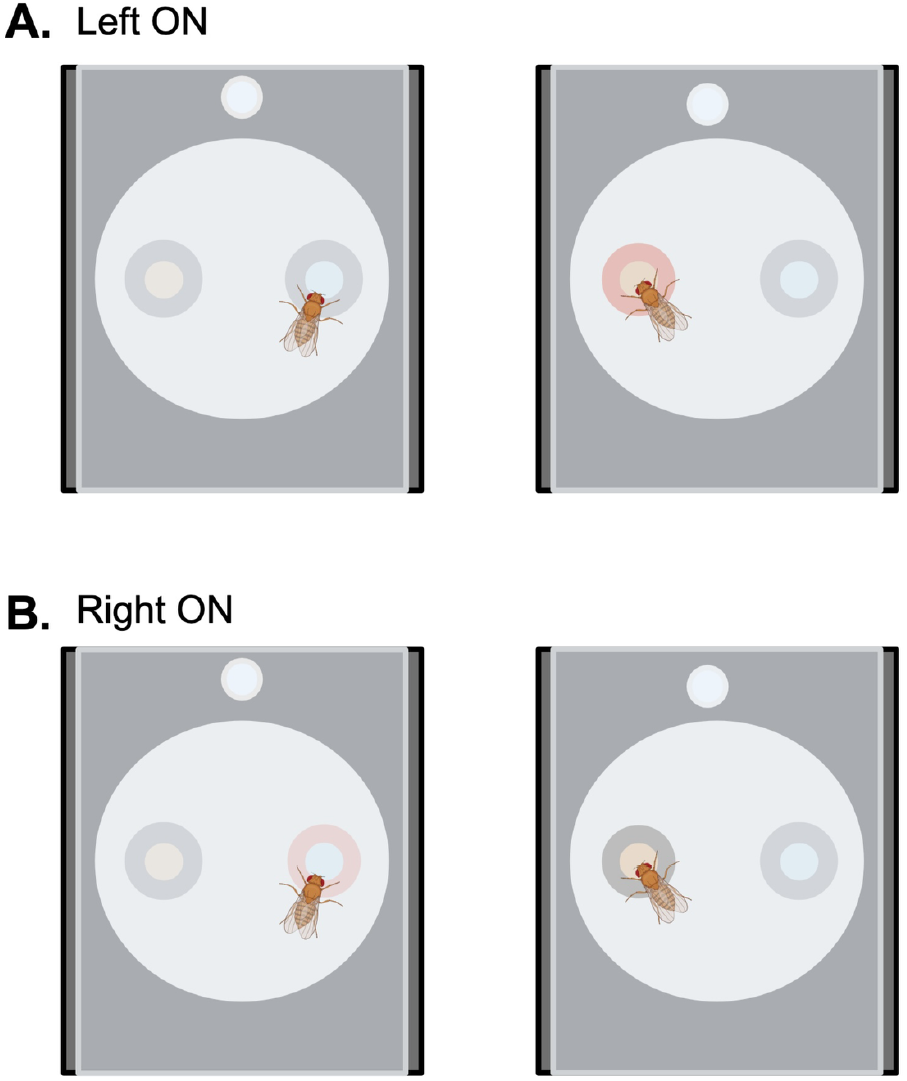
Closed-loop experiment pairing optogenetic neural manipulation with a specific food source. In closed-loop experiments, the LED is paired with the electrode associated with one designated food type. (A) When the LED is paired with the left electrode, interactions with the right food source do not trigger illumination. However, when the fly interacts with the left food source, the LED is activated to induce optogenetic neural manipulation. (B) Conversely, when the LED is paired with the right electrode, interactions with the right food source activate the LED to induce optogenetic neural manipulation, while interactions with the left food source do not trigger illumination.

**CRITICAL**: If an entire row is horizontal, this indicates a likely connection issue with that particular choice equipment and the rig. Please see troubleshooting section below.

7. Load flies into each arena using a fly aspirator.
  a. Ensure you load flies as per your experimental conditions.
  b. To load flies:
    i. Partially open each arena.
    ii. Tilt your fly aspirator such that the fly is pushed toward the back of the arena.
    iii. Immediately close chamber shut when flies are loaded.
8. Starting the experiment
  a. Hit “start” on the bonsai software to initiate the experiment.
  b. Set a lab timer for 50 minutes.
  c. Ensure the chambers are exposed to minimal to no lighting (darkness preferable)
  d. If you wish to run multiple experiments in the same day:
    i. Place the protein and sucrose food onto the mug warmer, set to 55□.
    ii. Alternate sides of presented food between experiments as a control.
9. Ending the experiment
  a. After 50 minutes, press “stop” in Bonsai to end the experiment.
  b. Re-collect the flies for additional tests or discard them based on institutional parameters.
10. Clean-up and resetting for another experiment
  a. To clean, use a sterile 10 μL pipette tip and insert it into each choice hole to push the food out.
  b. Use lightly damp lint-free tissue paper with distilled water to wipe up the residual food. You should also do this on the underside portion of the arenas.
  c. Once complete, ensure the arenas are dry using lint-free tissue paper and clear of all food residue.

### Behavior Protocol: OptoPAD Open-Loop assay

This section provides detailed methodology on performing a total consumption or choice experiment, where an experimenter wishes to determine the flies’ baseline consumption, or preference after neuronal activation or inhibition of food consumption between two different food types. This type of experiment is ideal for studying:

a. How activation or inhibition of specific cells impacts feeding behavior in real-time.

For example, you may wish to determine whether your particular fly line has a preference for protein or sucrose when a subset of neurons or glia cells are activated or inhibited with the LED module.

**TIMING:** 15-minute preparation, plus 50-minute experiment, with 15 minutes of cleanup/reset.

**CRITICAL:** Ensure flies have been raised on retinal food (1:500) or placed in a collection retinal vial (1:250) for 3 days prior to beginning the experiment.

1. Repeat steps 1 (PC preparation) through 6 (pre-experimental validation) above from the “flyPAD assay” protocol.
  a. Use either the total consumption or two-choice formats as desired.
2. When loading flies for optoPAD:
  a. You should include a non-LED control, or a subset of flies not exposed to light. See Figure 4 for an example experimental schematic.
  b. Example of groups:
    i. Non-LED - variable A level 1
    ii. Non-LED - variable A level 2
    iii. LED-variable A level 1
    iv. LED-variable A level 2
3. Situating the optogenetic modules for stimulation
  a. Place the provided optogenetic module(s) over each desired LED exposed arena.
  b. Turn the LED electronic module “ON”. You can set the voltage based on what percent LED stimulation you would like.
    i. EX. 3V: 50%
  c. Cover the non-LED groups using mousepads or similar to avoid any light saturation.

**CRITICAL SAFETY WARNING:** Do NOT look directly into the LED lights as this may cause visual impairment.

4. Repeat steps 8 (starting the experiment) through 10 (cleaning up) from the above flyPAD total consumption/two-choice protocol.

### Behavior Protocol: OptoPAD Closed-Loop Assay

In a closed-loop experiment, the fly’s own behavior is used as an input to modulate system output. For example, you may perform an experiment whereby every time a fly takes a sip of a particular substrate, there is consequential LED delivery, and real-time circuit manipulation. When synchronization between light and food consumption is achieved on the left side: the LED light turns on when the fly drinks from the left, and remains off on the right, and vice versa.

As opposed to open-loop, the closed-loop behavior is best to test:

a. If flies can learn to avoid or prefer a particular food option.
b. If neural activity alters decision-making at the moment of choice.

1. Pre-Experiment PC Preparation for “Left ON” experiment
  a. Open the Bonsai software.
  b. Select the OptoPAD program, “Octopad_B2_4_NewVersion_Blinking _v2”.
  c. When saving a directory:
    i. Create a new folder named “closedloop_date”.
    ii. In this folder, create two subfolder files, one “LEFT ON” and the second “RIGHT ON.”
    iii. Use naming conventions for conditions and flies per arena as described above.
2. Setting the LED to only trigger in the left channel
  a. Identify the node in Bonsai corresponding to the feeding-event stream from the left channel.
    i. Configure the workflow so that feeding events from the left channel are routed to the LED.
    ii. This will cause feeding interactions in the left channel to trigger LED activation.
  b. Identify the node corresponding to the feeding-event stream from the right channel.
    i. Configure the workflow so that feeding events from the right channel are recorded only and not connected to the LED digital output mode.
3. Repeat steps 2 (warming food) through 10 (cleaning up and resetting above).

**CRITICAL 1:** Wells should likely contain the same tastants, but this depends on experimental hypotheses and the research question.

4. Run “Right ON” experiments by following steps 1-3 above, except switching left and right channel instructions.

**NOTE:** It is recommended to run both “Left ON” and “Right ON” experiments for each condition you plan to test, and averaging results to account for any intrinsic side bias.

### Data Processing

Data will be processed using data analysis software provided by the manufacturer. Below, we will describe a workflow using MATLAB, but the same data handling and analytical concepts can be applied to other software. We will start from processing the raw data and conclude with recommended statistical applications.

1. Preparing the directory for data analytical software.
  a. Open the desired directory on the PC where you saved your data.
  b. There will be two data files and your.txt file.
    i. The larger file (∼58MB) is your data file from the 50 min experiment, and the one you need for analysis.
    ii. Ensure it is named appropriately as described above.
    iii. The smaller file (1-5MB) is your pre-experimental validation file. This data file needs to be deleted and/or removed from this directory.
2. Data Analytics
  a. Open your desired analytical software.
  b. Use manufacturer recommended analytical tools for each experimental protocol above.
    i. Analytical tools for various analytical software are provided by the manufacturer.
  c. Data will be saved in the original directory where the experiment was originally saved.
    i. Navigate to the experimental folder(s) you wish to analyze.
    ii. Open the “mrep” folder.
    iii. Open “DatainExcelFormat”.
3. Output metrics from the assay are described in Table 5.

**Table 5.** Data Output and Interpretation.

| Output | Description | Interpretation/Use |
| --- | --- | --- |
| Spill Quantity | Fraction of time of the experimental recording that the signal was saturated due to contact with the top electrode | Reflects food contacting electrodes due to improper loading or flies spilling the food during feeding. |
| Removed Non Eaters | Number (or proportion) of flies excluded because they did not meet a minimum feeding threshold | Flies that never fed or fed below a defined cutoff (2 or less activity bouts on both electrodes during the whole duration of the experiment). |
| Sip Durations | Duration (ms) of a single feeding contact | Proxy for ingestion strength or palatability. Together with Inter Sip Interval define the frequency of feeding (feeding rhythm) |
| Inter Sip Intervals (ISI) | Time between consecutive sips within a feeding episode | Short ISI = sustained feeding; long ISI = disengagement. |
| Number of Sips | Total count of feeding events per fly | Proxy for total intake |
| Activity Bout Number | Number of episodes of active exploration of food sources | Feeding initiation frequency |
| Activity Bout Duration | Median duration of active exploration of food sources | Sustained engagement with food |
| Feeding Bursts Number | Number of feeding bursts per fly. A feeding burst is a sub-behavior within an activity bout, characterized as rapid sequences of sips with very short ISIs | Micro-pattern of ingestion |
| Feeding Bursts Duration | Duration of each feeding burst | How intensely flies feed within bouts |
| Activity Bout IBI | Time between the end of one bout and the start of the next | Reflects satiety, motivation, or learning |
| Feeding Burst IBI | Pause between feeding bursts (within a bout). | Short feeding burst IBI's may be interpreted as increased motivation for a particular food source. |

**Table 6.**
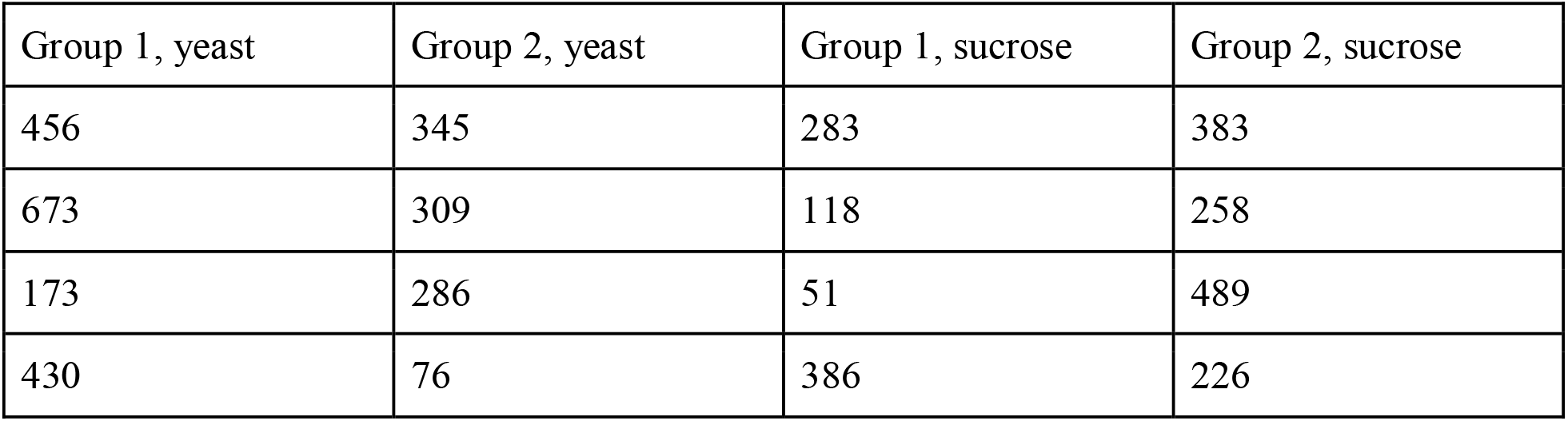
Number of Sips Example.

### Data Analysis: flyPAD or optoPAD (two-choice example)

Below are specific instructions for analyzing a flyPAD or optoPAD experiment where a fly was presented with a choice between either protein or sucrose. Please note that these are mock data used for instructional purposes and not actual data from any real experiments. In addition to looking at raw number of sips, below is a recommendation for creating a preference index.

### Preference Index

A preference index, as opposed to looking at raw number of sips, has several advantages. Due to several extraneous variables not limited to but including fly genetics, time of day, vial conditions, and/or housing parameters, there may be baseline differences in feeding behavior. A preference index normalizes the data to allow more interpretable comparisons between experiments.

A preference index will take the form of: 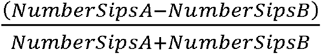

Such that:

1.0= Full preference for Substrate A

0.0 = No preference

-1.0 = Full preference for Substrate B

1. Calculating the Preference Index
  a. Open the “Number of Sips” panel within the excel document.
  b. For each group, apply the preference index formula above between your first and second substrate (Table 7).
    i. In the formula, substrate A is typically the substrate of interest.
2. Example for calculating a protein preference index:
  a. Subtract each sucrose value from the yeast value.
  b. Divide this by the addition of the total number of sips for both yeast and sucrose.
3. Statistical Applications
  a. Analysis Pipeline 1: Comparing the protein preference index values from group 1 and group 2 to examine group level differences in preference.
    i. Independent samples t-test (one group, two-levels).
      - Ex. Male vs female flies on protein preference.
    ii. Two-way ANOVA (two groups, two-levels)
      - Ex. Sex (male vs female) and starvation condition (fed vs starved) on protein preference.
  b. Analysis Pipeline 2: Comparing the preference value vs 0 to examine the absolute preference of each group.
    i. One-sample t-test against 0.
      - Ex. Despite group 1 (M = 0.10) having a significantly increased protein preference compared to group 2 (M = 0.0), both groups have no absolute preference (not significantly different from 0.0).
4. Other potentially useful calculated output metrics are featured in Table 8.

**Table 7.** Calculating a protein preference index.

| Group 1, protein (yeast) | Group 1, sucrose | Formula | Protein Preference Index Value |
| --- | --- | --- | --- |
| 456 | 283 | $(456-283) / (456 + 283)$ | 0.23 |
| 673 | 118 | ... | 0.70 |
| 173 | 51 | ... | 0.55 |
| 430 | 386 | ... | 0.04 |

**Table 8:** Calculated metrics from raw output, their description, and interpretation.

| Metric | Calculation | Description | Interpretation |
| --- | --- | --- | --- |
| Preference Index | $(SipA - SipB) / (SipA + SipB)$ | Relative feeding choice between two substrates | 1.0 = A preference<br>0.0 = No preference<br>-1.0 = B preference |
| Vigor | (SipA,B) /<br>Duration<br>Sip A,B | Measure of feeding<br>intensity (tempo) | Higher values indicate<br>stronger engagement with<br>the substrate, higher<br>urgency |
| Average sips per<br>activity bout | (SipA,B)<br>/ActivityBoutsA,B | Measure of feeding<br>continuity | Higher values indicate<br>longer, sustained<br>engagement with the<br>substrate |

### flyPAD and optoPAD Troubleshooting

1. Flies are not engaging with the assay or show very low feeding activity.
  a. Food has dried or solidified before or during the assay.
    i. Ensure assay food remains warm and liquid during loading and forms a dome-like shape in each well.
    ii. Low feeding motivation→ Increase dry starvation to 2–4 h prior to assay.
  b. Flies were exposed to CO□ within 24 h of the experiment.
    i. Avoid CO□ anesthesia within 24 h; transfer flies using a fly aspirator.
  c. Room temperature or humidity is outside the optimal range.
    i. Perform assays at 22–25 °C and ∼40–60% relative humidity.
2. The food dries or runs out during the assay.
  a. Low ambient humidity
    i. Ensure consistent (40-50%) room humidity.
  b. Extended preparation and loading food time before experiment start.
    i. Practice sufficiently to ensure this time is minimized. We recommend 10 minutes from loading the food to starting the experiment at maximum.
  c. Certain food substrates may dry faster than others.
    i. Load food in a consistent order and alternate sides between experiments.
  d. Experimental time is too long.
    i. The assay is optimally 30-50 minutes in length, only.
3. Flat/horizontal channels while starting assay.
  a. Food is touching the brass electrodes.
    i. Remove excess food with a piece of lab tissue.
    ii. Repeat the pre-test until all lines are wavy/zig-zag.
4. Entire row of channels is inactive.
  a. Loose or improperly connected flyPAD arena.
    i. Ensure the connection to the COM port is secure.
5. The LED lights are not functioning properly.
  a. LED module is not properly seated or there are faulty connections.
    i. Reposition the module to ensure LEDs are securely attached.
  b. Voltage box turned off or not properly set.
    i. Verify voltage output (3V or as desired) prior to starting the experiment.
6. Flies showing intrinsic bias for one probe/channel in a choice assay or closed-loop assay.
  a. Check if the chamber/arena is leveled. Adjust the length of screws at corners to adjust the level.
  b. Check if light is not evenly shed on the chamber in regular assay and if there is light bleed through in optogenetic assay.
  c. If bias persists, switch substances position in alternate trials and average the preference index of two back-to-back trials.
7. Issues with Data analysis, error messages.
  a. Files are incorrectly formatted or labeled (log file or bonsai file), Verify file naming matches the arena layout and experimental conditions.
  b. Ensure the log file contains the correct groups and substrates.
  c. Ensure the “test” file used to troubleshoot the capacitance detectors is deleted from the directory.
  d. Ensure your length of experiment time entered matches how long the behavior was recorded.

### Expected Outcomes

Feeding behavior in *Drosophila melanogaster* is a sensitive behavioral output that at baseline is dependent on sex, mating status, social conditioning, age, and genotype (e.g., Liu et al., 2024). Table 9 is a general table that outlines expected feeding outcomes across multiple variables.

**Table 9:** Expected Outcomes.

| Variable | Expected outcome on feeding behavior | References |
| --- | --- | --- |
| Age (young vs old) | Total feeding generally declines with age. Sucrose diet increases lifespan<sup>19-20</sup>. | Galenza et al., 2016<br>Wong et al., 2009 |
| Sex (male vs female) | Females (mated) generally consume more total food, and more protein, than males<sup>21-22</sup>. | Camus et al., 2018<br>Vargas et al., 2010 |
| Female mating status (virgin or mated) | Mated females generally have a higher preference for protein, and eat more, compared to virgins<sup>21,23</sup>. | Camus et al., 2018<br>Ribeiro & Dickson 2010 |
| Male mating status | Mixed results. Virgin males generally have an increased protein intake, which declines after ejaculation, and then is restored<sup>21,24</sup>. | Liu et al., 2024<br>Camus et al., 2018 |
| Nutritional History/Diet | Starvation or dietary restriction, and macronutrient specific diets affects total feeding,choice behavior, fecunditiy, and fitness<sup>24-26</sup>. | Liu et al., 2024<br>Itskov & Rubeiro, 2013<br>Lee et al., 2008 |
| Temperature | At lower temperatures, *Drosophila melanogaster* prefers plant foods as opposed to yeast<sup>27</sup>. | Brankatschk et al. 2018 |
| Genetic Background | Common control lines (*Canton-S*, *W118*, *Oregon-R*) show baseline differences in feeding behavior<sup>20-21,26</sup>. | Camus et al. 2018<br>Wong et al. 2009<br>Lee et al. 2008 |

## Acknowledgements

We would like to thank Tarandeep Singh Dadyala, Sumaya Noor Smithy, and Laura White for helpful discussions and feedback on experimental design and protocol optimization. We would also like to thank the manufacturer, Pavel Itskov, for providing technical details when we were writing the manuscript. This work is supported by a Maximizing Investigators’ Research Award to L.S. (National Institute of General Medical Sciences R35GM147504).

## Author Contributions

All authors were involved with conceptualizing, planning, and reviewing all versions of the manuscript. N.J.C: Primary writer and editor for all versions of the manuscript.

M.E.: Figures, resource tables, reviewing all versions of the manuscript.

I.T.S.: Figures, resource tables, writing-closed-loop protocol, reviewing all versions of the manuscript. L.S.: Supervision, funding acquisition, writing – reviewed, edited, and provided feedback on all versions of the manuscript. All authors have read and approved the final version of the manuscript.

## Data Availability

All data provided in this manuscript are mock datasets for educational and instructional purposes. For inquiries regarding the data analysis software and associated tools used with the flyPAD/optoPAD system, readers should consult the manufacturer documentation provided with the equipment, or contact the corresponding authors for further information.

